# Predicting Disease-Specific Histone Modifications and Functional Effects of Non-coding Variants by Leveraging DNA Language Models

**DOI:** 10.1101/2025.06.15.659749

**Authors:** Xiaoyu Wang, Tong Pan, Sihan Chen, Geoffrey I. Webb, Yunzhe Jiang, Joel Rozowsky, Mark Gerstein, Jiangning Song

**Affiliations:** Monash Biomedicine Discovery Institute, Monash University, Melbourne, VIC 3800, Australia; Monash AI Institute, Monash University, Melbourne, VIC 3800, Australia; Department of Data Science and Artificial Intelligence, Monash University, Melbourne, VIC 3800, Australia; Program in Computational Biology and Bioinformatics, Yale University, New Haven, CT 06520, USA; Department of Molecular Biophysics and Biochemistry, Yale University, New Haven, CT 06520, USA; Department of Computer Science, Yale University, New Haven, CT 06520, USA; Department of Statistics and Data Science, Yale University, New Haven, CT 06520, USA; Department of Biomedical Informatics and Data Science, Yale University, New Haven, CT 06520, USA

**Keywords:** Histone modification, Alzheimer’s disease, Deep learning, Language model

## Abstract

**Background:** Epigenetic modifications play a vital role in the pathogenesis of human diseases, particularly neurodegenerative disorders such as Alzheimer’s disease (AD), where dysregulated histone modifications are strongly implicated in disease mechanisms. While recent advances underscore the importance of accurately identifying these modifications to elucidate their contribution to AD pathology, existing computational methods remain limited by their generic approaches that overlook disease-specific epigenetic signatures.

**Results:** To bridge this gap, we developed a novel large language model (LLM)-based deep learning framework tailored for disease-contextual prediction of histone modifications and variant effects. Focusing on AD as a case study, we integrated epigenomic data from multiple patient samples to construct a comprehensive, disease-specific histone modification dataset, enabling our model to learn AD-associated molecular signatures. A key innovation of our approach is the incorporation of a Mixture of Experts (MoE) architecture, which effectively distinguishes between disease and healthy epigenetic states, allowing for precise identification of AD-relevant epigenetic modification patterns. Our model demonstrates robust performance in disease-specific histone modification prediction, achieving mean area under receiver-operating curves (AUROCs) ranging from 0.7863 to 0.9142, significantly outperforming existing state-of-the-art methods that lack disease context. Beyond accurate modification site prediction, our framework provides important biological insights by successfully prioritizing AD-associated genetic variants, which show significant enrichment in disease-relevant pathways, supporting their potential functional role in AD pathogenesis. These findings suggest that differential modification loci identified by our model may represent key regulatory elements in AD.

**Conclusions:** Our framework establishes a powerful new paradigm for epigenetic research that can be extended to other complex diseases, offering both a valuable tool for variant effect interpretation and a promising strategy for uncovering novel disease mechanisms through epigenetic profiling.

## Background

Epigenetic mechanisms, encompassing histone modifications and DNA methylation, play pivotal roles in gene regulation and disease pathogenesis. Post-translational histone modifications, particularly H3K27 acetylation (H3K27ac), trimethylation (H3K27me3) and H3K4 trimethylation (H3K4me3), serve as critical determinants of chromatin state and transcriptional regulation. Histone modification can be determined experimentally by Chromatin immunoprecipitation followed by sequencing (Chip-seq). Notably, accumulating research indicates that epigenetic dysregulation contributes substantially to neurodegenerative pathogenesis, such as Alzheimer’s disease (AD) [1] and Parkinson’s disease (PD) [2].

Over the last decade, numerous studies have explored histone modification prediction, and existing computational approaches for epigenetic prediction primarily fall into two categories: (i) Variant effect prediction models (e.g., DeepSEA [3], ExPecto [4] and Sei [5]) that predict the effects of sequence variants across multiple chromatin features; (ii) Histone-specific predictors (e.g., DeepHistone [6], Histone-Net [7] and DeepPTM [8]) designed exclusively for histone modification site prediction. The differences in histone modifications exist not only across different cell types but also among various diseases. However, research on disease-associated histone modification patterns prediction remains limited, hindering our understanding of their pathogenic mechanisms.

Two fundamental challenges thus emerge: (1) how to effectively learn disease-specific epigenetic modifications under sample size constraints; and (2) how to computationally model the disease-contextual regulatory grammar to gain insights into the molecular mechanism underlying disease. To overcome the limitations of existing methods, we introduce the EpiModX, a new large language model (LLM)-based deep learning framework specifically designed for disease-contextual prediction of histone modification and variant effects. Recent advances in large-scale DNA language models such as DNABERT [9], Nucleotide Transformer [10], and Caduceus [11], enable comprehensive learning of genomic grammatical structures, syntactic patterns, and semantic relationships directly from sequence data. Traditional methods for analyzing epigenetic modifications have relied primarily on raw DNA sequences without fully considering the complex “grammar” that governs how genes are regulated. These approaches miss important higher-level patterns in how different parts of the genome work together to control gene activity. To overcome these limitations, our new framework makes two key advances. First, we use a DNA language model that can understand the hidden rules of genomic organization, including how distant DNA segments interact and how their meaning changes in different contexts. This allows us to extract meaningful patterns even when working with limited epigenetic data from individual patients. Second, we combine this DNA language model with a specialized Mixture of Experts (MoE) module that can simultaneously recognize both general rules of gene regulation and disease-specific contexts. This dual approach is particularly important because diseases like atherosclerosis [12] and Alzheimer’s disease [1], show distinct patterns of histone modifications, likely due to changes in how regulatory enzymes are recruited to specific DNA sequences. The MoE component allows our model to capture disease-specific epigenetic variations while still maintaining an understanding of fundamental regulatory principles, thereby providing a more complete picture of how epigenetic changes contribute to different health conditions.

In this work, Alzheimer’s disease is used as a case study to validate the feasibility of EpiModX. Alzheimer’s disease, a prototypical age-associated neurodegenerative disorder with strong heritable components, currently affects 12% of the worldwide population beyond age 65 [13]. Alzheimer’s disease progression from early-stage mild cognitive impairment to late-stage dementia affects tens of millions of individuals worldwide, representing a growing global health challenge. While several variants in AD-related genes (e.g., *APP*, *PSEN1*, and *PSEN2*) have been identified in early-onset Alzheimer’s disease (EOAD), these account for merely 1-5% of total AD cases [14]. In contrast, late-onset AD (LOAD), which constitutes the majority of cases, demonstrates substantially more complex and less predictable genetic risk patterns. Substantial evidence demonstrates significant associations between altered histone modification patterns and Alzheimer’s disease pathogenesis. Nativio et al. found that H3K27ac and H3K9ac are enriched in Alzheimer’s Disease, and the increased global H3K27ac/H3K9ac in an AD fly model worsened Aβ42-mediated neurotoxicity [1]. Histone acetylome-wide association study identified the 4,162 differential H3K27ac peaks, which were enriched in Alzheimer’s disease-related biological pathways [15]. Therefore, understanding how epigenetic alterations contribute to disease pathogenesis is crucial for elucidating Alzheimer’s disease mechanisms [16].

While numerous studies have identified disease-associated epigenetic changes, a critical unresolved challenge has been determining whether SNPs within histone modification peaks functionally influence these epigenetic marks. Existing approaches have primarily focused on whether SNPs overlap with histone modification regions [1], which cannot distinguish causal variants from coincidental co-locations. Our framework overcomes this limitation by predicting how genetic variants directly affect histone modifications across individuals of various disease states and infer the functional impact of variants. This capability enables mechanistic investigation of disease origins, even in studies with limited sample sizes, by revealing which SNPs actively alter the epigenetic landscape rather than merely residing near modified regions.

The current study makes three significant advances: (1) We develop a powerful deep learning model that achieves state-of-the-art performance in predicting histone modification sites by incorporating large language models to understand DNA’s regulatory grammar; (2) We create the first framework capable of predicting disease-specific histone modification patterns, moving beyond generic epigenetic predictions; and (3) We establish a new approach for identifying functionally relevant non-coding variants in Alzheimer’s disease by analyzing their differential epigenetic effects. Although we demonstrate our model using Alzheimer’s disease as a case study, the framework is readily applicable to other diseases with epigenetic components. The system can analyze any genetic variant of interest, enabling researchers to investigate how histone modification patterns diverge between health and disease states, and uncover the potential mechanisms driving these differences. This represents a powerful new paradigm for understanding the epigenetic basis of complex disorders and accelerating therapeutic discovery.

## Results

### Overview of deep learning framework

To develop our disease-specific prediction model, we constructed a comprehensive disease-specific histone modification dataset using ENCODE consortium data from individuals across the Alzheimer’s disease spectrum, including those with no cognitive impairment (NCI), mild cognitive impairment (MCI), cognitive impairment (CI), Alzheimer’s disease (AD), and both AD with cognitive impairment (AD+CI). This dataset incorporated ChIP-seq profiles for three key histone marks—H3K4me3, H3K27ac, and H3K27me3—across these clinical groups (all sample accession IDs provided in **Supplementary Table 1**). Our genome-wide analysis identified 417,343 H3K27ac, 105,964 H3K4me3, and 261,468 H3K27me3 peaks across individuals. We merged peak intervals across individuals based on genomic coordinates and overlap coverage percentage (see **Methods**), while randomly selecting an equal number of negative control regions (excluding ENCODE blacklist regions) that showed no overlap with called peaks. Notably, the majority of peaks were under 4,000 bp in length, with fewer than 3% exceeding 6,000 bp across all modifications and disease groups (**Supplementary Figure 1**), consistent with typical chromatin feature distributions.

Building on this dataset, we developed EpiModX, a new deep learning framework that combines a DNA language model with a (MoE) architecture (**Figure 1**) to predict disease-specific histone modifications. Our framework employs multi-task learning where all prediction tasks share a common fine-tuned language model backbone (Caduceus [11]), enabling knowledge transfer while maintaining task-specific adaptation through dedicated MoE modules. Caduceus, built on the long-range Mamba architecture and trained via masked language modeling (MLM) on the human genome, processes DNA sequences as input to generate contextual embeddings that capture complex genomic information. Drawing from experimental evidence showing disease-associated alterations in epigenetic modifier activity and binding motif mutations studies [1], we designed our model to systematically learn these genome-wide context-specific epigenetic patterns. Following Chen et al.’s approach [17], we implemented customised MoE module to capture disease group-specific histone modification patterns, with additional parameters allocated for each clinical group’s distinctive features. A routing network was employed to select the Top-K experts for each group, while our optimization strategy combines a loss function that maximizes inter-group mutual information and parameter divergence to enhance disease-specific pattern recognition, with minimization of cross-entropy loss for individual prediction tasks. This allows the model to effectively distinguish disease-specific epigenetic characteristics while maintaining awareness of universal genetic patterns captured by the foundational DNA language model.

**Figure 1.**
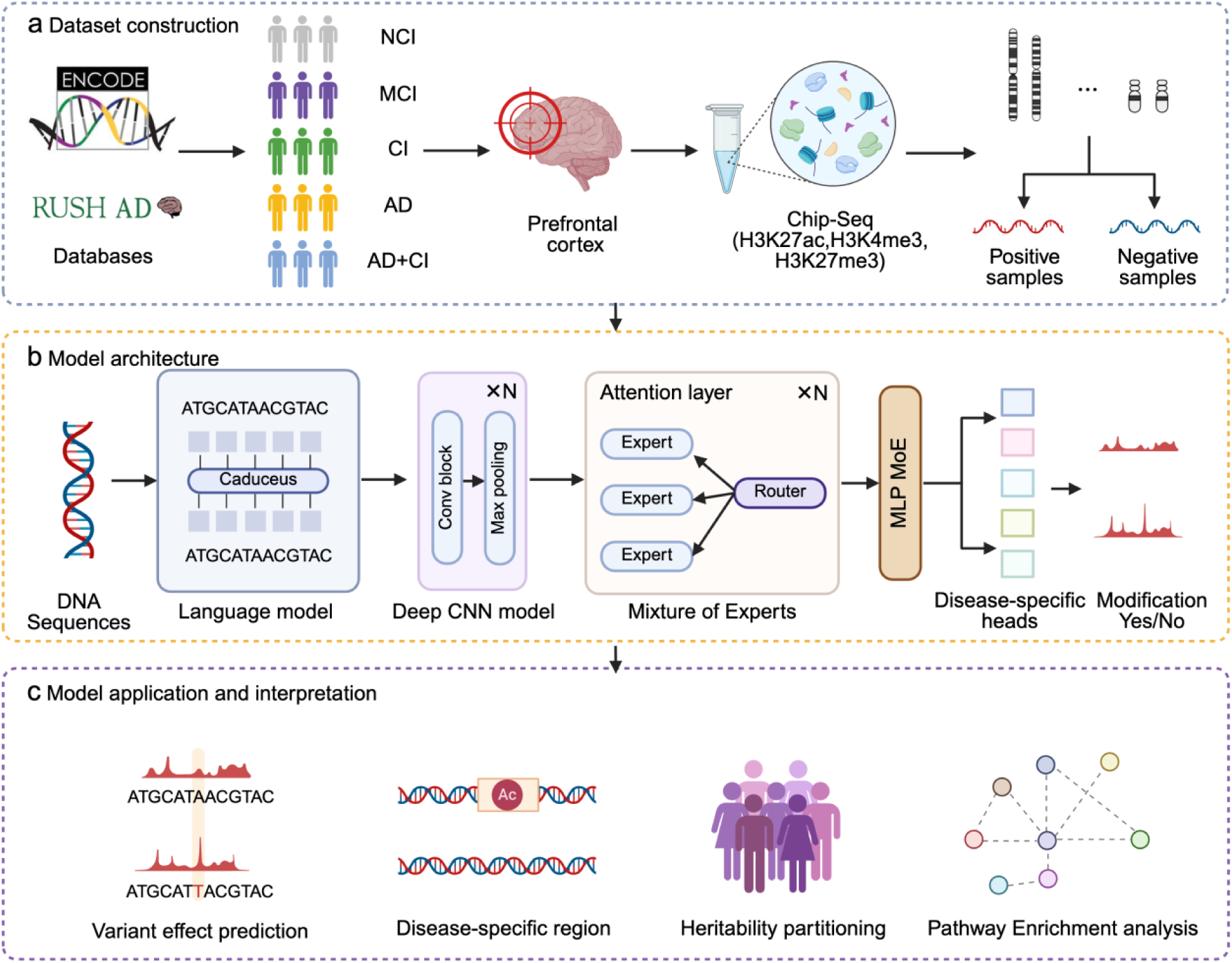
Overview of the EpiModX framework. (a) Data construction pipeline showing sample acquisition and processing from five clinical groups: no cognitive impairment (NCI), mild cognitive impairment (MCI), cognitive impairment (CI), Alzheimer’s disease (AD), and AD with cognitive impairment (AD+CI). (b) Model architecture integrating three key components: DNA language model, convolutional neural networks, and Mixture of Experts (MoE) modules. (c) Downstream model applications include variant effect prediction, disease-specific analysis, and mechanistic interpretation.

### Benchmarking disease-specific histone modification prediction

Our model demonstrates robust capability in predicting histone modification sites directly from DNA sequences, effectively capturing cis-regulatory patterns. For comprehensive evaluation, we assessed the model’s performance on independent test datasets using four key metrics: Accuracy (ACC), F1-score (F1), Area Under the Receiver Operating Characteristic Curve (AUROC), and Area Under the Precision-Recall Curve (AUPRC). Using a chromosome-wise data partition strategy, we designated chromosome 10 for validation and chromosomes 8 and 9 for testing, ensuring rigorous evaluation of generalization capability.

Comparative analysis against state-of-the-art modification predictors (DeepSEA, Expecto, and DeepHistone) revealed our model’s superior performance across all histone marks. Specifically, our method achieved average AUROCs of 0.8104 (H3K27ac), 0.9117 (H3K4me3) and 0.7863 (H3K27me3), representing statistically significant improvements (*P*<0.001, two-sided Wilcoxon signed rank test) of 1.72%, 2.12%, and 4.74%, respectively, over existing methods (**Figure 2d and Supplementary Tables 2-3**). The AUROC and AUPRC curves (**Supplementary Figure 2**) further confirm our model’s enhanced predictive power. Two key innovations drive this performance advantage: (1) A whole genome-scale DNA language model that is capable of capturing long-range epigenetic dependencies through whole-genome contextual sequence learning, and (2) A specialized MoE architecture that effectively discriminates disease-specific regulatory patterns from control contexts. This dual approach enables superior cross-condition prediction performance compared to existing methods.

**Figure 2.**
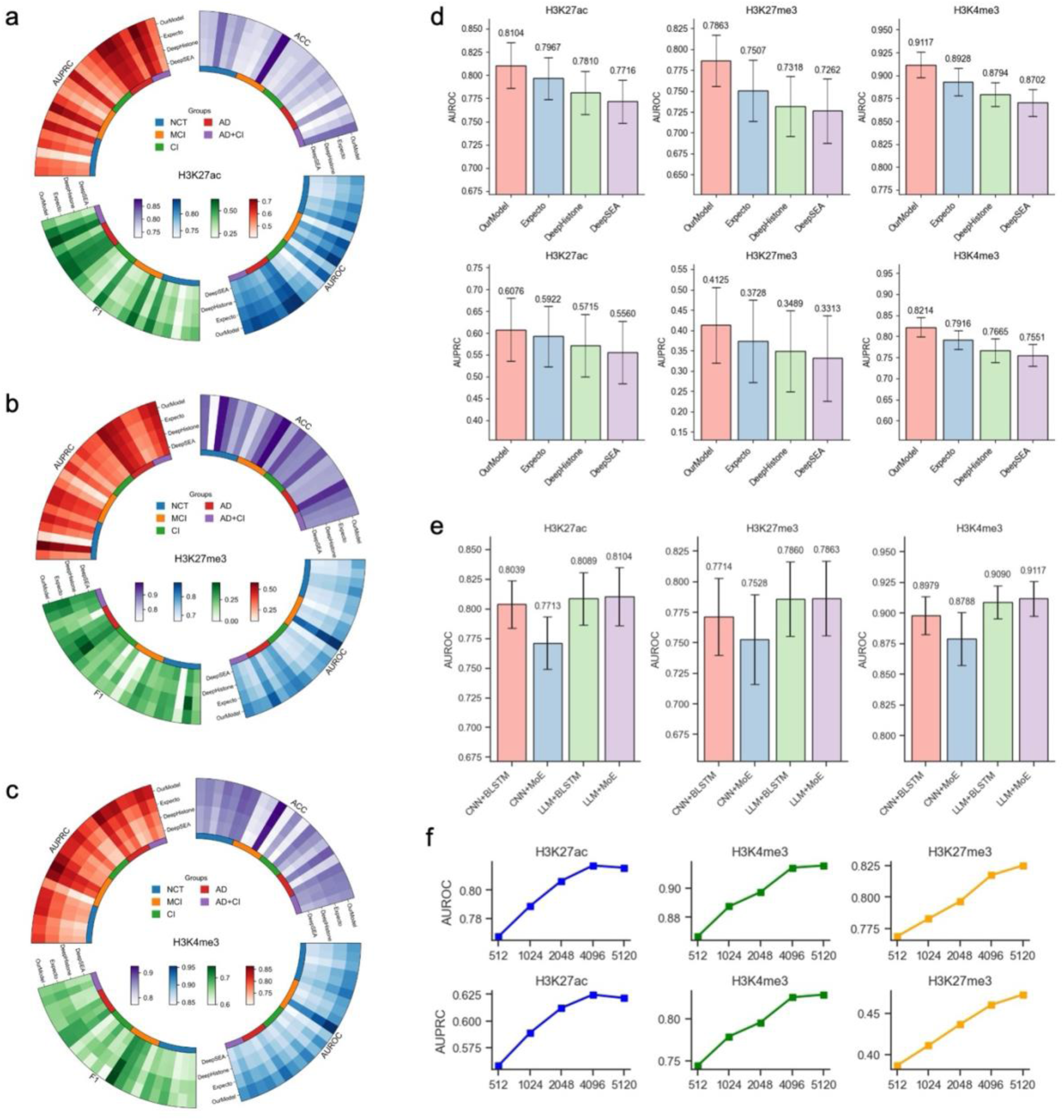
Model performance evaluation and ablation studies. Individual-level performance for (a) H3K27ac, (b) H3K27me3, and (c) H3K4me3 modifications. (d) Performance comparison of our model with state-of-the-art methods (DeepSEA, ExPecto, DeepHistone). (e) Ablation analysis of model components: convolution neural networks (CNN); bidirectional long short-term memory (BLSTM); Mixture of Experts (MoE). (f) Performance variation across multiple input sequence lengths (512-5,120 bp), with optimal performance achieved at 4,096 bp.

At the individual sample level, our model achieved consistently strong performance despite substantial inter-individual epigenetic variability in epigenetic modification patterns (**Figures 2a-c**). Across all evaluation metrics (AUROC, AUPRC, F1 and ACC), our framework outperforms existing methods for every test sample, demonstrating both robust generalization and reliable prediction accuracy for individual cases. This individual-level consistency, combined with overall superior performance, highlights our model’s potential for precision epigenomics applications.

### Model ablation analysis and model optimization

To assess the contributions of each component in our framework, we performed ablation studies focusing on two core modules: the DNA language model (LLM) and the MoE module. We compared our full LLM-MoE model against three alternative architectures: (1) CNN+BLSTM, replacing both key modules with conventional deep learning components; (2) CNN+MoE, retaining only the MoE module; and (3) LLM+BLSTM, preserving just the language model. The comparative analysis of our model variants (**Figure 2e** and **Supplementary Table 4-5**) demonstrated the superior performance of the complete LLM-MoE architecture across all histone modification datasets. The language model component proved especially crucial, as the model incorporating the LLM consistently outperformed their CNN-based counterparts, likely due to the LLM’s ability to capture complex genomic grammar and long-range dependencies in DNA sequences that are essential for accurate epigenetic prediction. This was evidenced by a substantial performance drop when using the conventional CNN architectures, which failed to maintain the necessary genomic sequence context for the MoE module to function effectively. Similarly, replacing the MoE module with BLSTM led to reduced performance, highlighting the importance of the MoE’s specialized design for discriminating disease-specific epigenetic patterns. These ablation results collectively demonstrate that both the DNA language model and MoE architecture make unique and complementary contributions to our framework’s performance, with the LLM providing foundational genomic understanding and the MoE enabling fine-grained disease context modeling.

The length of DNA sequences used as input plays a crucial role in accurately predicting epigenetic modifications. Through systematic evaluation of input sequence lengths ranging from 512 bp to 5,120 bp (**Figure 2f** and **Supplementary Table 6**), we determined that 4096 bp represents the optimal balance between predictive performance and computational efficiency. This length consistently outperformed shorter sequences in both AUROC and AUPRC, capturing sufficient genomic context to identify modification patterns while remaining computationally practical. Interestingly, for H3K27ac modifications specifically, we found that expanding the sequence window beyond 4096 bp actually reduced the predictive performance, likely because excessive flanking DNA sequence introduces noise without providing additional relevant regulatory information. The 4,096 bp window appears to optimally encompass both the core modification sites and their surrounding regulatory elements, making it the most effective choice for our predictive model while maintaining reasonable computational demands.

### The model successfully captured disease-specific epigenetic signatures

To assess our model’s ability to capture disease-specific epigenetic contexts, we performed a cross-disease group analysis using the precision and recall metrics, which offer more subtle insights into prediction biases across disease groups than traditional AUROC and AUPRC measures. Statistical evaluation using the two-sided Wilcoxon rank-sum test revealed significant inter-group performance differences between the NCI and AD groups for H3K27ac sites (*P*-value<=0.02) (**Figure 3a-b**. The AD-optimized model exhibited lower precision but higher recall when predicting acetylation sites in the NCI group, indicating a biologically meaningful bias when AD-derived sequences were more frequently classified as acetylation sites compared to those NCI-derived sequences. This finding aligns with established multi-omics evidence showing expanded H3K27ac occupancy in AD pathogenesis [1], suggesting that our model successfully recapitulates known disease-associated epigenetic expansion. Parallel analyses of H3K4me3 modifications showed similar inter-group (between the NCI and AD groups) differences (**Supplementary Figure 3**), while H3K27me3 patterns remained stable across disease states, consistent with its role as a repressive mark less susceptible to disease-associated fluctuations. To gain deeper insights into the disease-specific patterns learned by our model, we analyzed and visualized the weight scores across different groups (**Figure 3c**). Our analysis revealed distinct motif preferences and epigenetic signatures characteristic of each disease state. The model demonstrated differential weighting of transcription factor binding sites and other regulatory elements between groups. These differential weighting patterns not only validate our model’s ability to discern disease-specific regulatory grammars but also highlight potential mechanistic drivers of epigenetic dysregulation in disease progression.

**Figure 3.**
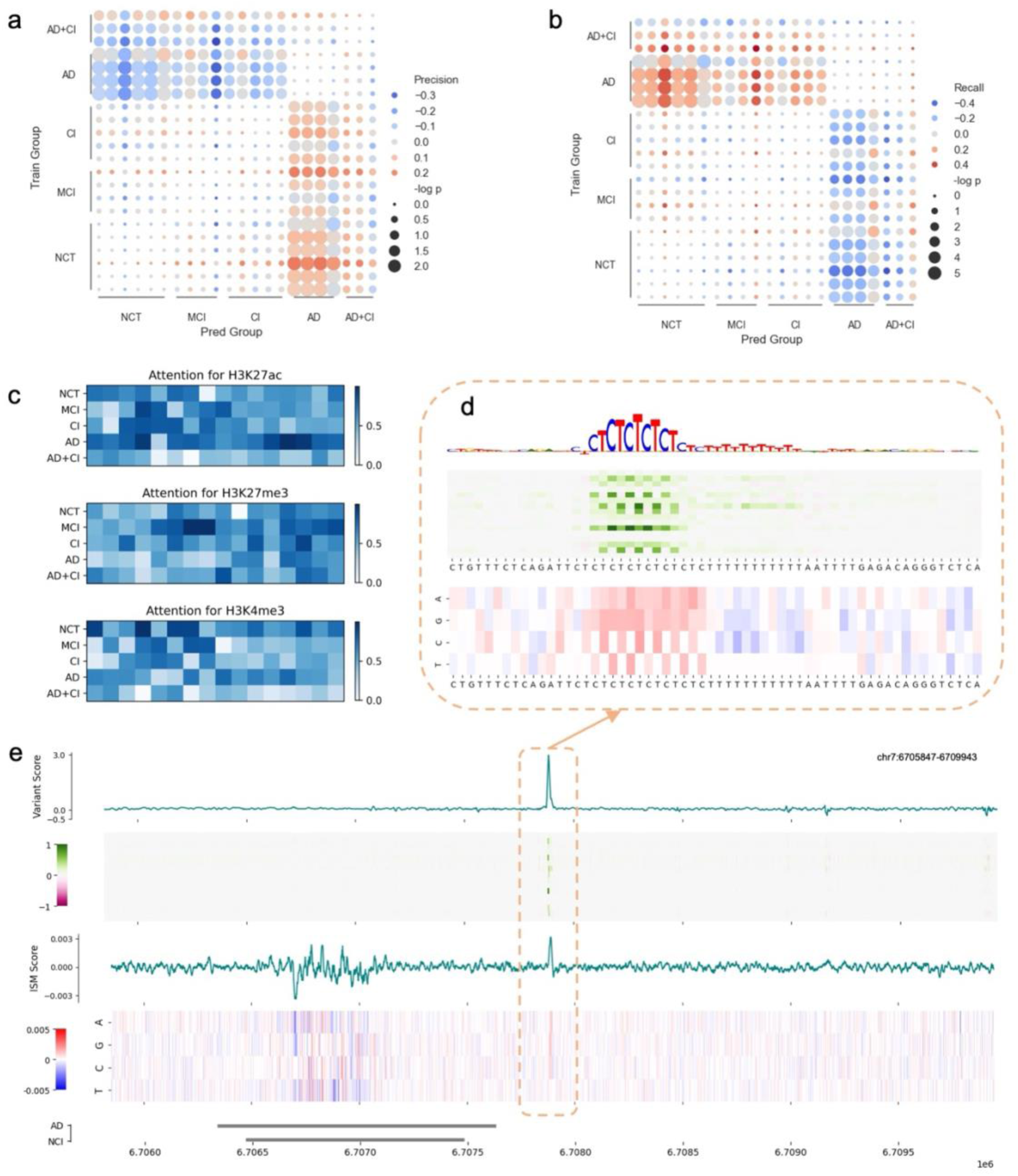
Cross-disease group analysis of H3K27ac patterns. (a) Precision and (b) recall metrics across disease groups, where dot size represents −*log*_10_ P value and color gradient indicates performance deviation in cross-disease analysis. (c) Disease-specific attention weights across experts for different disease groups, visualized as heatmaps. (d) Differential contribution region identified by Gradient × Input analysis. We visualise the contribution score and the (e) In silico mutagenesis (ISM) scores for a representative differential H3K27ac site between NCI and AD groups, with corresponding peak tracks shown in grey.

To elucidate the molecular mechanisms underlying disease-associated histone modifications, we implemented two complementary approaches to quantify sequence contribution scores: in silico saturation mutagenesis and Gradient × input. Using a representative genomic locus (chr7:6705847-6709943) as a case study, we systematically evaluated the functional impact of all possible nucleotide substitutions across this region (**Figure 3e**) through in silico saturation mutagenesis. This approach could precisely identify modification-sensitive sites despite the absence of explicit positional information in our training data, revealing critical cis-regulatory elements through systematic sequence perturbation analysis (**Figure 3e**).

The Gradient × input method provides complementary insights by quantifying the contribution of individual nucleotides to model predictions. This alternative approach is particularly useful for identifying disease-specific regulatory patterns, as evidenced by the differential contribution scores (defined as ΔGradient × input) calculated between the AD and NCI groups. We identified and visualized the motif (**Figure 3d)** that might underlie the differential modifications using TFMoDisco [18]. Notably, several predicted differential modification sites were found to co-localize with short tandem repeats, aligning with emerging evidence that such repetitive sequences could contribute to Alzheimer’s disease risk through epigenetic mechanisms [19]. Short tandem repeats have been suggested to influence the binding affinities between DNA and transcription factors, potentially contributing to gene expression [20].

### haQTL analysis of variants predicted by the DL model and validation using GWAS data

A critical challenge in epigenetic research involves identifying genetic variants that functionally influence histone acetylation quantitative trait loci (haQTLs) [21] without requiring large-scale experimental profiling. To validate our model’s capability to identify haQTLs, we constructed a comprehensive haQTL benchmark dataset using 5,161 brain-specific haQTLs from published studies [22] as positive samples. Negative controls were carefully selected from three genomic distance intervals (2kb, 5kb and 10kb downstream of known haQTLs) to ensure comparable genomic context while excluding true regulatory variants. This stratified approach allowed us to systematically evaluate our model’s precision in distinguishing functional from non-functional variants.

The analysis revealed significantly higher importance scores for true haQTLs compared to all variants in negative control regions (2kb downstream: *P* = 3.32 × 10^-39^; 5kb downstream: *P* = 2.75 × 10^-102^, 10kb downstream: *P*= 9.23 ×10^-101^; One-sided Wilcoxon rank-sum test) (**Figure 4a**), demonstrating our model’s ability to distinguish functional regulatory variants from non-functional ones. Notably, the relatively higher importance scores observed for variants located 2 kb downstream may be attributed to the presence of potential haQTLs. In contrast, 5kb and 10kb downstream controls were indistinguishable (*P* = 0.36, One-sided Wilcoxon rank-sum test), establishing a clear boundary for functional prediction. To comprehensively evaluate our model’s performance in haQTL identification, we calculated the AUROCs and AUPRCs based on the haQTLs and variants located in the downstream regions of 2 kb, 5 kb and 10 kb, respectively (**Figure 4b**). Our model showed good performance across all relative distances to their targets.

**Figure 4.**
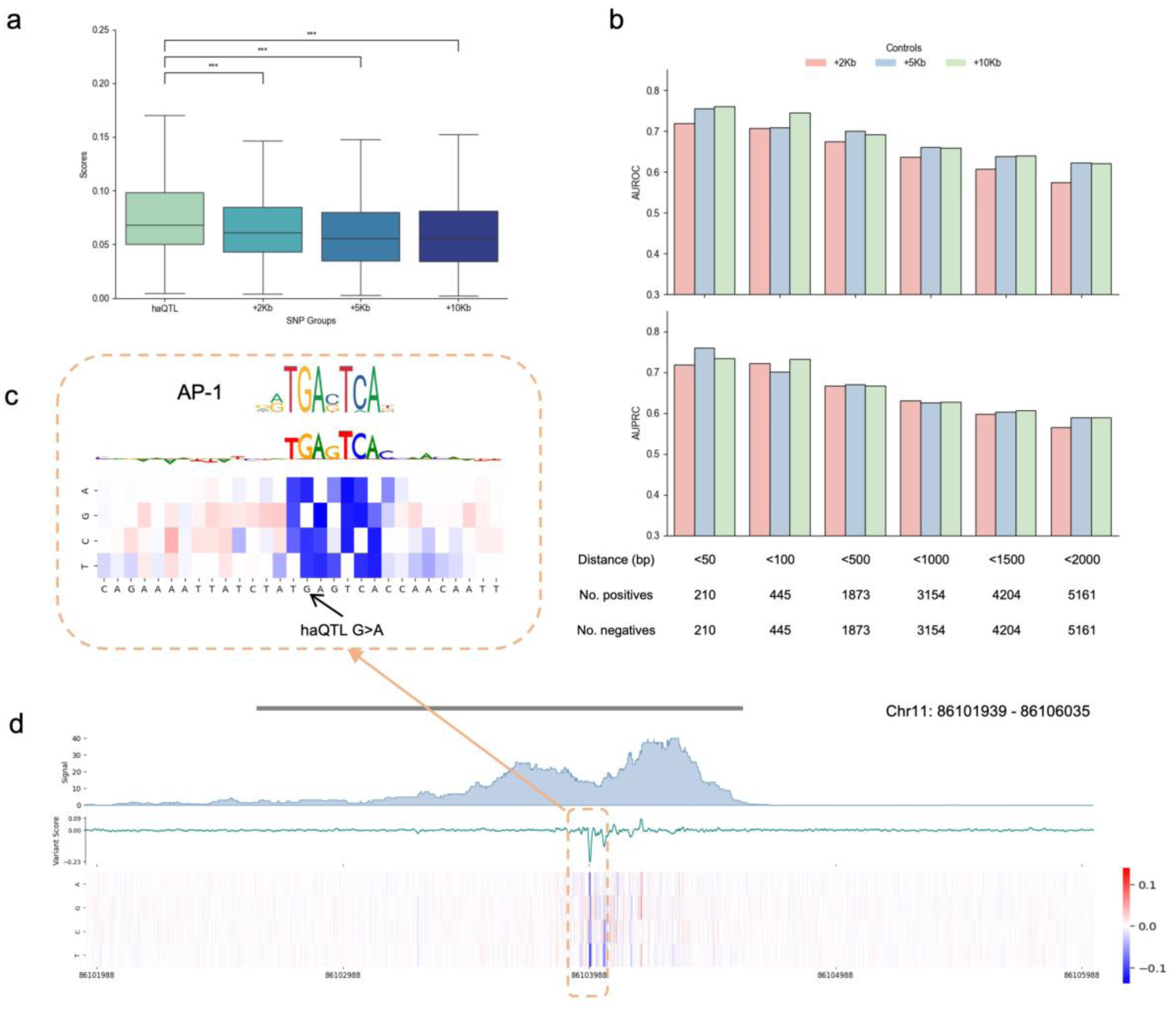
haQTL prediction and validation. (a) Importance score distribution for validated haQTLs versus downstream control variants (2kb/5kb/10kb). Boxplots measure importance scores. (b) AUROCs and AUPRCs for haQTLs predicted by the H3K27ac model across all relative distances to targets. (c-d) In silico mutagenesis (ISM) analysis of a brain-specific haQTL (Chr11:86,103,988) and its 4kb genomic region (Chr11: 86,101,939-86,106,035) containing this haQTL, revealing disruption of an AP-1 transcriptional factor binding motif associated with H3K27ac regulation.

A compelling example of our model’s biological interpretability comes from its identification of the chr10:86103988 variant, a known haQTL identified through GWAS studies. Our model not only accurately predicted this variant’s regulatory effect but also revealed its precise molecular mechanism. Through in silico mutagenesis of the surrounding genomic region, we observed that this variant significantly alters an AP-1 transcription factor binding motif (**Figure 4c-d**) [23]. This finding has particular biological relevance as AP-1 is a well-established regulator of H3K27ac dynamics in neuronal development and function [24]. The variant’s location within this key regulatory motif suggests it may influence AD pathogenesis by modulating the recruitment of epigenetic writers/readers to neuronal enhancers. This case exemplifies how our model can contribute beyond simple variant identification to provide testable hypotheses about the underlying functional mechanisms.

### SNP prioritization and partitioning heritability in disease

Our model also provides a powerful approach for prioritising disease-related SNPs by integrating epigenetic information with genetic association data. Given that histone modification peak regions can span up to 6,000 bp (**Figure S1**), simply identifying SNP-overlapping regions offers limited biological insight. To evaluate our model’s SNP prioritization capability, we analyzed Alzheimer’s disease GWAS summary statistics from Kunkle et al. [25] using stratified linkage disequilibrium (LD) score regression (S-LDSC) [26]. This approach allowed us to quantify the heritability enrichment in our predicted functional regions, thereby extending beyond traditional methods that only consider physical overlap between SNPs and histone marks. The results revealed significant enrichment of disease-associated variants in our predicted differential H3K27ac sites (Enrichment = 10.500, *p*-value = 0.0014) (**Figure 5b**), highlighting our model’s ability to identify biologically meaningful relationships between genetic variation and epigenetic regulation.

**Figure 5.**
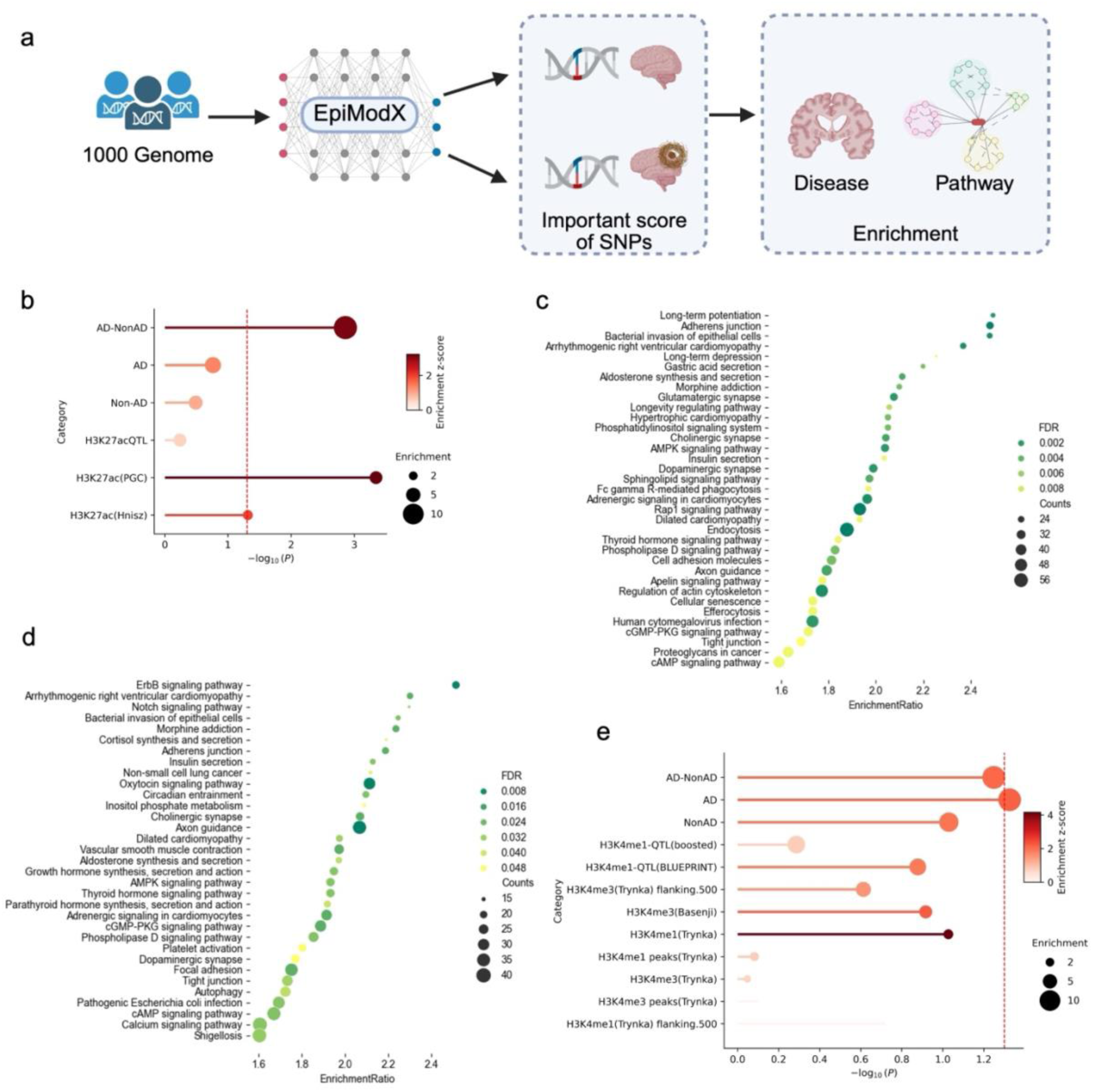
Disease pathway enrichment and heritability analysis. (a) Framework for predicting disease-specific variant effects, quantified by changes in the probability scores across different groups. (b) Genome-wide heritability partitioning using predicted H3K27ac annotations versus established epigenetic annotations. (c) KEGG pathways enriched in differential H3K27ac sites (FDR<0.001). (d) Pathway enrichment for AD-specific H3K4me3 predictions (FDR<0.01). (e) Genome-wide heritability partitioning for H3K4me3- and H3K4me1-related annotations.

When comparing our deep learning-predicted epigenetic annotations with established baseline annotations (including H3K27ac QTL, PGC and Hnisz datasets) [27], we observed superior performance in capturing AD-associated variants. The differential analysis between NCI and AD groups revealed that SNPs with high differential effects were specifically enriched in AD-relevant genomic regions, while predictions from either group alone failed to show significant enrichment.

In addition to H3K27ac, our model’s performance also extended to H3K4me3 modifications, where AD-specific contribution scores exhibited significant enrichment (Enrichment = 10.034, *P*-value = 0.0472) compared to other H3K4me3-related annotations (**Figure 5e**). This consistency across different histone marks underscores our model’s generalizability and precision in effectively capturing nucleotide-level contributions to disease-associated epigenetic regulation. The model’s capacity to learn these disease-specific epigenetic patterns from limited training data suggests it could be particularly valuable for studying other complex disorders where large epigenetic datasets are unavailable. Furthermore, our model’s accurate prediction of variant effects provides a powerful tool for prioritizing non-coding mutations in genetic studies and identifying potential therapeutic targets in regulatory elements.

### Predicted risk variants demonstrate pathway enrichment specific to Alzheimer’s disease pathology

Our pathway analysis of disease-specific variants predicted by the H3K27ac and H3K4me3 models revealed significant biological insights into Alzheimer’s disease pathogenesis. Consistent with the literature showing minimal changes in H3K4me3 patterns in patients with AD [28], we focused our analysis on more dynamic H3K27ac and H3K4me3 modifications. Using variants from the 1000 Genomes Project Phase 3 and annotated by the Ensembl’s Variant Effect Predictor [29], we identified significantly enriched pathways through KEGG pathway enrichment analysis using WebGestalt [30]. The H3K27ac model predictions identified 35 significantly enriched pathways (FDR *P*-value<=0.001) strongly associated with AD mechanisms (**Figure 5c and Supplementary Table 7)**, including synaptic plasticity pathways (long-term potentiation [31] and long-term depression [32]), cell junction integrity (adherent junction [33] and cell adhesion molecules [34]), and pathways related to gut-brain-microbiome interactions (bacterial invasion of epithelial cells [35] and gastric acid secretion [36]). These findings suggest that disease-associated alterations in histone acetylation patterns may stem from modified histone acetyltransferase (HAT) activity in AD, potentially through altered binding affinities or recognition of motif patterns, ultimately leading to dysregulation of critical neuronal and cellular pathways. Moreover, a recent study has shown that HAT activators may function as promising therapeutic modulator in the treatment of Alzheimer’s disease [37].

The H3K4me3 model analysis similarly identified 32 significantly enriched pathways (FDR *P*-value<=0.05), with notable overlap in cardiac pathways (arrhythmogenic right ventricular cardiomyopathy [38]) and infection-related processes (bacterial invasion of epithelial cells [35]), while also revealing unique neural pathways including ErbB signaling [39], Notch signaling [40], oxytocin signaling [41], and axon guidance [42] (**Figure 5d and Supplementary Table 8)**. The convergence of certain pathways across both histone marks suggests coordinated epigenetic dysregulation in AD, while mark-specific pathways likely reflect distinct aspects of disease pathogenesis. Of particular interest is the identification of developmental pathways, such as Notch signaling and axon guidance, which may indicate reactivation of developmental programs in neurodegeneration. Overall, the pathway enrichment analyses not only validate the biological relevance of our model’s predictions but also provide new mechanistic insights into how epigenetic dysregulation at specific genomic loci may contribute to AD through perturbation of key molecular networks involved in neuronal function, cellular homeostasis, and potentially even microbial interactions.

## Discussion

A critical challenge in genomic medicine involves deciphering the functional consequences of non-coding variants and their epigenetic effects in disease contexts. While GWAS studies have successfully identified numerous disease-associated loci, these population-level approaches require massive genomic-scale sample sizes and typically lack mechanistic insights. Existing computational methods for epigenetic prediction, including DeepSEA and Expecto, face similar limitations—while effective for general chromatin feature prediction, they fail to incorporate disease-specific contextual information, thereby significantly restricting their utility for precision medicine applications.

To bridge this gap, we developed a novel deep learning framework that synergistically combines a DNA language model with an MoE architecture to capture disease-specific histone modification patterns using limited samples. Our approach addresses two fundamental challenges: (1) the desirable need to model disease-specific epigenetic patterns from limited patient samples, and (2) the requirement for interpretable predictions that link genomic variants to functional consequences. By constructing a comprehensive AD-specific dataset encompassing multiple disease stages (NCI, MCI, AD, and AD+CI), we demonstrated that our model’s superior performance over existing methods, achieving state-of-the-art prediction accuracy for key histone modifications. The success of our framework stems from its dual-capacity architecture: the DNA language model effectively captures long-range genomic dependencies and latent sequence grammars, while the MoE module specializes in recognizing disease-contextual epigenetic patterns.

Our model’s performance was rigorously validated through multiple downstream tasks that demonstrate its ability to capture biologically meaningful disease-specific patterns. In predicting haQTLs, the model showed excellent discriminative power, with true haQTLs exhibiting significantly higher contribution scores than background variants. This capability was particularly relevant in our analysis of brain-specific H3K27ac haQTLs, where the model successfully distinguished functional regulatory variants from non-functional neighboring SNPs while maintaining high accuracy. The model’s disease relevance was further confirmed through S-LDSC using Alzheimer’s disease GWAS summary statistics, which revealed significant heritability enrichment in our predicted variant sets compared to existing annotations. To elucidate the biological pathways associated with the predicted H3K27ac and H3K4me3 variants, we performed KEGG pathway analysis. The results identified several AD-relevant pathways, including synaptic plasticity pathways, cell junction integrity and infection-related processes, all of which have established roles in AD pathogenesis. These findings not only validate our model’s biological relevance but also provide novel insights into how epigenetic dysregulation in these pathways may contribute to disease progression. Importantly, by integrating nucleotide-level variant effect prediction with pathway-level functional analysis, our framework provides a powerful new approach for prioritizing disease-relevant non-coding variants, revealing their mechanistic roles, and generating testable hypotheses about epigenetic contributions to disease pathogenesis. Notably, our method identifies key disease-related regulatory elements, such as short tandem repeats, which have been suggested as novel therapeutic targets for Huntington’s disease [43, 44].

While our model demonstrates robust performance in predicting disease-associated histone modifications and analyzing genetic heritability, several limitations should be acknowledged. First, the current framework does not incorporate individual genetic variants due to dataset constraints, potentially missing opportunities to explore how specific mutations impact epigenetic regulation. Second, while our model while our model successfully identifies disease-specific patterns, the training dataset’s limited sample size may affect generalizability across diverse populations and disease stages. Another important consideration is the resolution of our analysis, which operates at the bulk tissue level rather than single-cell resolution. This limitation prevents the detection of cell-type-specific regulatory mechanisms that could provide more precise insights into disease pathogenesis.

Several promising directions exist for further improvement and application. First, expanding the training dataset with additional samples would enhance the model’s predictive power and provide more comprehensive coverage of epigenetic modification patterns. Although our approach successfully operates with limited data, increased sample sizes would improve statistical power and enable the detection of more subtle disease-associated epigenetic changes. Second, experimental validation of our computational predictions represents a crucial step forward. The model’s ability to identify differential modification sites and prioritize functional variants offers a valuable resource for guiding targeted wet-lab studies, potentially accelerating the discovery of novel epigenetic targets for Alzheimer’s disease intervention. Third, the generalizability of our framework suggests broad applicability across diverse disease contexts. The methodology can be potentially applied to other conditions with significant epigenetic components, including various cancer types [45] and Parkinson’s disease [46], where it may help elucidate disease mechanisms and identify therapeutic targets. Future adaptations could incorporate cell-type specific information and additional epigenetic marks to create even more comprehensive models of disease-associated regulatory changes.

### Conclusions

In this study, we developed a new LLM-based deep learning framework termed EpiModX for predicting disease-specific histone modifications, with particular application to Alzheimer’s disease. Our model demonstrates superior performance compared to existing state-of-the-art methods, as evidenced by comprehensive benchmarking across multiple evaluation metrics. Through systematic ablation studies, we validated the contributions of each component, particularly highlighting the importance of the DNA language model for capturing genomic context and the MoE architecture for discerning disease-specific epigenetic patterns. The model’s capabilities extend beyond accurate modification site prediction to include haQTL identification and functional variant analysis, providing a powerful tool for epigenetic research. Notably, our predicted epigenetic annotations show significant enrichment for AD-associated variants compared to other conventional modification annotations, establishing a stronger link between histone modification patterns and disease pathology. These findings suggest that genome-wide alterations in histone modification coverage may contribute to AD pathogenesis through dysregulation of gene expression networks. The ability of our framework to pinpoint differential modification sites and analyze variant effects opens new avenues for understanding the epigenetic basis of complex diseases and identifying potential therapeutic targets. Future applications of this approach could provide deeper insights into how epigenetic dysregulation contributes to neurodegenerative diseases and other pathological conditions.

## Materials and methods

### Data collection

We obtained all ChIP-seq data for histone modifications (H3K27ac, H3K4me3 and H3K27me3) from the Rush Alzheimer’s Disease Study through the ENCODE database, focusing on dorsolateral prefrontal cortex samples through uniform processing pipelines. To ensure data quality, we excluded regions from ENCODE blacklists and trimmed sequences containing unresolved bases. For dataset integration, we merged overlapping sequences (>80% overlap) and used the combined intervals, centering each on the peak middle point and extending to 4,096bp to capture relevant regulatory contexts. Positive samples were defined as sequences containing either: (1) at least one peak in the central 2 kb region, or >50% of sequence length identified as histone modification sites, with exclusion of peaks with lengths exceeding size thresholds. Negative samples were randomly sampled from GRCh38/hg38 human genomic regions devoid of histone modifications, matched for length and genomic context. All genomic coordinates and analyses were based on the GRCh38/hg38 reference genome. The complete list of epigenetic profiles and sample details are provided in **Supplementary Table 1**.

### Model implementation details

Our framework employs a hybrid architecture that processes Chip-seq profile peak calls (binary annotations) to predict continuous probability values for histone modification likelihood. The model integrates three core components: (1) a large DNA language model for sequence embedding, (2) deep convolutional networks for feature extraction, and (3) customized Mixture of Experts (MoE) modules for disease-specific pattern recognition. We implemented the model using PyTorch, with full architectural details available in our GitHub code repository.

For sequence encoding, we utilized Caduceus [11], a pretrained large DNA language model, which was implemented based on the Mamba architecture [47]. Using the Caduceus model, we encoded each DNA sequence into a 16-dimensional numerical feature vector. The model was initialized with pretrained weights retrieved from HuggingFace. To reduce spatial dimensions and aggregate information, we adopted a deep neural network module, which contains 4 convolutional neural network layers (kernel size is 5). Max pooling layers were inserted after every convolutional layer.

To identify the difference between different groups, we adopted a method called mixture of experts, in this model. Inspired by previous work, we develop the MoE method based on the Mod-Squad methods. The MoE module contains two MoE attention blocks and MoE MLP blocks. We customised the MoE attention block to capture the disease-specific information. The assumption is that the motifs or other recognize patterns of the disease groups are different, while our MoE module could combine the disease-specific patterns to predict the histone modification. Each MoE block contains 16 experts along with five disease-specific routing networks, corresponding to five disease groups. The output of MoE attention layer is formulated as follows [17]:

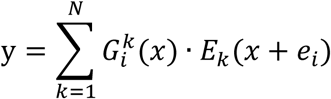

where *i* is the disease group, G^k^_i_ is the Noisy Routing network, and *e*_*i*_ denotes the disease-specific embedding. *E*_*k*_ represents the expert, which is the attention head.

Finally, the model architecture also incorporates five specialized output heads, one for each disease group.

### Model training and evaluation

The main model was implemented in Pytorch and it was trained on NVIDIA A100 with the batch size of 8. In this study, we adopt the AdamW optimizer and the learning rate is 5 × 10^−5^. An early stopping strategy was employed to prevent overfitting, with the stopping criterion set at 5 epochs. The loss function is the sum of binary cross-entropy loss and the loss of MoE module, which maximize mutual information between the experts and disease-specific groups. The combined loss is defined as:

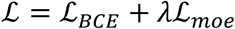

where λ is the weight for MoE layer loss, which is 0.5 in this study. The combined loss enhances the model’s capability to discriminate between disease groups.

We split the dataset into training, validation, and testing sets based on the chromosome. Chromosome 10 was used as the validation set, and chromosomes 8 and 9 were used as the test set. The performance comparison between our model and other state-of-the-art methods was conducted on the same test dataset. The large DNA language model was fine-tuned using pretrained weights obtained from Hugging Face.

### Model interpretability

To provide some explanation for this model, Captum package [48] was used to calculate the important scores. The importance scores are defined as Gradient × input, while the differential importance scores are defined as:

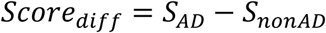

where the *S*_*AD*_ is the average important score of the disease group, calculated based on the embeddings generated by fine-tuned DNA language models. AD and AD+CI groups are combined to form a general AD cohort (N=7), while the NCI group are the non-AD cohort (N=6).

The silico mutagenesis scores were calculated as the difference in model output probabilities between reference and mutated sequences. Using in silico mutagenesis at single-nucleotide resolution, we computed position-wise contribution scores and constructed a 4 × *L* importance matrix (where L represents sequence length), which was subsequently visualized as a heatmap. To identify key regulatory sites, we derived an aggregated importance score by summing ISM values across a 5-bp sliding window.

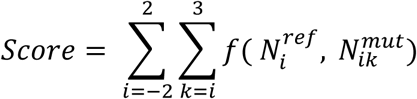

where the *k* is represents the alt alleles. We calculate the scores from position −2 to +2 relative to the central reference point, capturing a ±2 bp window to assess local mutational or epigenetic signal patterns.

### haQTL prediction

We obtained a set of haQTL-gARE pairs using Genome-wide association studies (GWAS) of brain tissue from the study of Hou et al.[22]. The haQTLs within 2 kb of the target region are selected in this study. To predict the variant effect scores with our model, we used in-silico mutagenesis (ISM) to calculate the importance score of each variant. For each haQTL, we compute its importance score by summing the importance scores of the 5-bp sequence centered on the variant, in which the nucleotides are replaced with random bases. The variant effects of haQTLs are calculated based on the change in predicted probability of the NCI group.

### Enrichment analysis

For enrichment analysis, we computed the importance scores for all 1000 Genomes Project Phase 3 variants at 10bp resolution, with genomic coordinates lifted from hg19 to hg38 using the UCSC LiftOver tool. Differential importance scores were defined as novel annotations for disease-specific analysis, where the predicted disease-associated variants with the highest scores were prioritized for downstream evaluation. The top 1000 disease-associated genes identified through this scoring system were subjected to KEGG pathway enrichment analysis using a false discovery rate (FDR) threshold of 0.05 and default parameters for other settings.

### Heritability partitioning

For disease-specific epigenetic analysis, we implemented a differential modification score (|ΔScore|) as a primary annotation metric, where the threshold was set at the 95th percentile of absolute values genome-wide. SNPs exceeding this threshold (top 5% by magnitude) were assigned a value of 1, while all others were coded as 0. We evaluated these predictions using stratified LD score regression (S-LDSC) with baseline-LD model v2.3 annotations, focusing particularly on histone-related annotations. Enrichment *P*-values were derived from the S-LDSC output scores. All heritability partitioning analyses were performed using the standard S-LDSC implementation available at https://github.com/bulik/ldsc.

## Data availability and resources

All datasets generated and analysed during this study, along with the EpiModX implementation, are publicly available in the EpiModX GitHub repository (https://github.com/yanwu20/EpiModX). For benchmarking purposes, comparison model architectures were obtained from their respective GitHub repositories. Human population variant data for heritability partitioning were sourced from the 1000 Genomes Project Phase 3, while baseline-LD model annotations (v2.3) were obtained from https://zenodo.org/records/10515792. Alzheimer’s disease GWAS summary statistics [25] were downloaded from the GWAS Catalog (accession GCST007511) at https://www.ebi.ac.uk/gwas/.

## Authors’ contributions

X.W developed the algorithm, conducted the experiments, and wrote the first draft of the manuscript. P.T and S.C performed the data analysis and contributed to figure preparation and data visualization. Y.J, J.R, M.G, G.W and J.S revised the manuscript and provided critical feedback. All authors critically reviewed and approved the final manuscript.

## References

1. Nativio R, Lan Y, Donahue G, Sidoli S, Berson A, Srinivasan AR, Shcherbakova O, Amlie-Wolf A, Nie J, Cui X: An integrated multi-omics approach identifies epigenetic alterations associated with Alzheimer’s disease. Nature genetics 2020, 52:1024–1035.

2. Gordevicius J, Li P, Marshall LL, Killinger BA, Lang S, Ensink E, Kuhn NC, Cui W, Maroof N, Lauria R, et al: Epigenetic inactivation of the autophagy–lysosomal system in appendix in Parkinson’s disease. Nature Communications 2021, 12:5134.

3. Zhou J, Troyanskaya OG: Predicting effects of noncoding variants with deep learning– based sequence model. Nature Methods 2015, 12:931–934.

4. Zhou J, Theesfeld CL, Yao K, Chen KM, Wong AK, Troyanskaya OG: Deep learning sequence-based ab initio prediction of variant effects on expression and disease risk. Nature Genetics 2018, 50:1171–1179.

5. Chen KM, Wong AK, Troyanskaya OG, Zhou J: A sequence-based global map of regulatory activity for deciphering human genetics. Nature Genetics 2022, 54:940–949.

6. Yin Q, Wu M, Liu Q, Lv H, Jiang R: DeepHistone: a deep learning approach to predicting histone modifications. BMC Genomics 2019, 20:193.

7. Asim MN, Ibrahim MA, Malik MI, Razzak I, Dengel A, Ahmed S: Histone-net: a multi-paradigm computational framework for histone occupancy and modification prediction. Complex & Intelligent Systems 2023, 9:399–419.

8. Baisya DR, Lonardi S: Prediction of histone post-translational modifications using deep learning. Bioinformatics 2020, 36:5610–5617.

9. Ji Y, Zhou Z, Liu H, Davuluri RV: DNABERT: pre-trained Bidirectional Encoder Representations from Transformers model for DNA-language in genome. Bioinformatics 2021, 37:2112–2120.

10. Dalla-Torre H, Gonzalez L, Mendoza-Revilla J, Lopez Carranza N, Grzywaczewski AH, Oteri F, Dallago C, Trop E, de Almeida BP, Sirelkhatim H, et al: Nucleotide Transformer: building and evaluating robust foundation models for human genomics. Nature Methods 2025, 22:287–297.

11. Schiff Y, Kao C-H, Gokaslan A, Dao T, Gu A, Kuleshov V: Caduceus: Bi-directional equivariant long-range dna sequence modeling. arXiv preprint arXiv:240303234 2024.

12. Lin Y, Qiu T, Wei G, Que Y, Wang W, Kong Y, Xie T, Chen X: Role of Histone Post-Translational Modifications in Inflammatory Diseases. Front Immunol 2022, 13:852272.

13. Matthews KA, Xu W, Gaglioti AH, Holt JB, Croft JB, Mack D, McGuire LC: Racial and ethnic estimates of Alzheimer’s disease and related dementias in the United States (2015–2060) in adults aged≥ 65 years. Alzheimer’s & Dementia 2019, 15:17–24.

14. Reitz C, Brayne C, Mayeux R: Epidemiology of Alzheimer disease. Nature Reviews Neurology 2011, 7:137–152.

15. Marzi SJ, Leung SK, Ribarska T, Hannon E, Smith AR, Pishva E, Poschmann J, Moore K, Troakes C, Al-Sarraj S, et al: A histone acetylome-wide association study of Alzheimer’s disease identifies disease-associated H3K27ac differences in the entorhinal cortex. Nature Neuroscience 2018, 21:1618–1627.

16. Heneka MT, van der Flier WM, Jessen F, Hoozemanns J, Thal DR, Boche D, Brosseron F, Teunissen C, Zetterberg H, Jacobs AH, et al: Neuroinflammation in Alzheimer disease. Nature Reviews Immunology 2024.

17. Chen Z, Shen Y, Ding M, Chen Z, Zhao H, Learned-Miller EG, Gan C: Mod-squad: Designing mixtures of experts as modular multi-task learners. In Proceedings of the IEEE/CVF Conference on Computer Vision and Pattern Recognition. 2023: 11828–11837.

18. Shrikumar A, Tian K, Avsec Ž, Shcherbina A, Banerjee A, Sharmin M, Nair S, Kundaje A: Technical note on transcription factor motif discovery from importance scores (TF-MoDISco) version 0.5. 6.5. arXiv preprint arXiv:181100416 2018.

19. Guo MH, Lee W-P, Vardarajan B, Schellenberg GD, Phillips-Cremins JE: Polygenic burden of short tandem repeat expansions promotes risk for Alzheimer’s disease. Nature Communications 2025, 16:1126.

20. Horton CA, Alexandari AM, Hayes MGB, Marklund E, Schaepe JM, Aditham AK, Shah N, Suzuki PH, Shrikumar A, Afek A, et al: Short tandem repeats bind transcription factors to tune eukaryotic gene expression. Science 2023, 381:eadd1250.

21. Pelikan RC, Kelly JA, Fu Y, Lareau CA, Tessneer KL, Wiley GB, Wiley MM, Glenn SB, Harley JB, Guthridge JM, et al: Enhancer histone-QTLs are enriched on autoimmune risk haplotypes and influence gene expression within chromatin networks. Nature Communications 2018, 9:2905.

22. Hou L, Xiong X, Park Y, Boix C, James B, Sun N, He L, Patel A, Zhang Z, Molinie B, et al: Multitissue H3K27ac profiling of GTEx samples links epigenomic variation to disease. Nature Genetics 2023, 55:1665–1676.

23. Rauluseviciute I, Riudavets-Puig R, Blanc-Mathieu R, Castro-Mondragon Jaime A, Ferenc K, Kumar V, Lemma RB, Lucas J, Chèneby J, Baranasic D, et al: JASPAR 2024: 20th anniversary of the open-access database of transcription factor binding profiles. Nucleic Acids Research 2023, 52:D174–D182.

24. Malik AN, Vierbuchen T, Hemberg M, Rubin AA, Ling E, Couch CH, Stroud H, Spiegel I, Farh KK-H, Harmin DA, Greenberg ME: Genome-wide identification and characterization of functional neuronal activity–dependent enhancers. Nature Neuroscience 2014, 17:1330–1339.

25. Kunkle BW, Grenier-Boley B, Sims R, Bis JC, Damotte V, Naj AC, Boland A, Vronskaya M, van der Lee SJ, Amlie-Wolf A, et al: Genetic meta-analysis of diagnosed Alzheimer’s disease identifies new risk loci and implicates Aβ, tau, immunity and lipid processing. Nature Genetics 2019, 51:414–430.

26. Finucane HK, Bulik-Sullivan B, Gusev A, Trynka G, Reshef Y, Loh P-R, Anttila V, Xu H, Zang C, Farh K, et al: Partitioning heritability by functional annotation using genome-wide association summary statistics. Nature Genetics 2015, 47:1228–1235.

27. Dey KK, van de Geijn B, Kim SS, Hormozdiari F, Kelley DR, Price AL: Evaluating the informativeness of deep learning annotations for human complex diseases. Nature Communications 2020, 11:4703.

28. Cao Q, Wang W, Williams JB, Yang F, Wang Z-J, Yan Z: Targeting histone K4 trimethylation for treatment of cognitive and synaptic deficits in mouse models of Alzheimer’s disease. Science Advances 2020, 6:eabc8096.

29. McLaren W, Gil L, Hunt SE, Riat HS, Ritchie GRS, Thormann A, Flicek P, Cunningham F: The Ensembl Variant Effect Predictor. Genome Biology 2016, 17:122.

30. Elizarraras JM, Liao Y, Shi Z, Zhu Q, Pico Alexander R, Zhang B: WebGestalt 2024: faster gene set analysis and new support for metabolomics and multi-omics. Nucleic Acids Research 2024, 52:W415–W421.

31. Wang T, Zhou YQ, Wang Y, Zhang L, Zhu X, Wang XY, Wang JH, Han LK, Meng J, Zhang X, et al: Long-term potentiation-based screening identifies neuronal PYGM as a synaptic plasticity regulator participating in Alzheimer’s disease. Zool Res 2023, 44:867–881.

32. Penney J, Tsai L-H: Histone deacetylases in memory and cognition. Science Signaling 2014, 7:re12-re12.

33. Sweeney MD, Sagare AP, Zlokovic BV: Blood–brain barrier breakdown in Alzheimer disease and other neurodegenerative disorders. Nature Reviews Neurology 2018, 14:133–150.

34. Hu J, Lin SL, Schachner M: A fragment of cell adhesion molecule L1 reduces amyloid-β plaques in a mouse model of Alzheimer’s disease. Cell Death & Disease 2022, 13:48.

35. Wu S-C, Cao Z-S, Chang K-M, Juang J-L: Intestinal microbial dysbiosis aggravates the progression of Alzheimer’s disease in Drosophila. Nature Communications 2017, 8:24.

36. Park A-M, Tsunoda I: Helicobacter pylori infection in the stomach induces neuroinflammation: the potential roles of bacterial outer membrane vesicles in an animal model of Alzheimer’s disease. Inflammation and Regeneration 2022, 42:39.

37. Bhatnagar A, Thomas CM, Nge GG, Zaya A, Dasari R, Chongtham N, Manandhar B, Kortagere S, Elefant F: Tip60 HAT activators as therapeutic modulators for Alzheimer’s disease. Nature Communications 2025, 16:3347.

38. Saeed A, Lopez O, Cohen A, Reis SE: Cardiovascular Disease and Alzheimer’s Disease: The Heart–Brain Axis. Journal of the American Heart Association 2023, 12:e030780.

39. Deng X-h, Liu X-y, Wei Y-h, Wang K, Zhu J-r, Zhong J-j, Zheng J-y, Guo R, Zhu Y-f, Ye Q-h, et al: ErbB4 deficiency exacerbates olfactory dysfunction in an early-stage Alzheimer’s disease mouse model. Acta Pharmacologica Sinica 2024, 45:2497–2512.

40. Kapoor A, Nation DA: Role of Notch signaling in neurovascular aging and Alzheimer’s disease. In Seminars in cell & developmental biology. Elsevier; 2021: 90–97.

41. Selles MC, Fortuna JTS, de Faria YPR, Siqueira LD, Lima-Filho R, Longo BM, Froemke RC, Chao MV, Ferreira ST: Oxytocin attenuates microglial activation and restores social and non-social memory in APP/PS1 Alzheimer model mice. iScience 2023, 26.

42. Zhang L, Qi Z, Li J, Li M, Du X, Wang S, Zhou G, Xu B, Liu W, Xi S: Roles and mechanisms of axon-guidance molecules in Alzheimer’s disease. Molecular neurobiology 2021, 58:3290–3307.

43. Matuszek Z, Arbab M, Kesavan M, Hsu A, Roy JCL, Zhao J, Yu T, Weisburd B, Newby GA, Doherty NJ, et al: Base editing of trinucleotide repeats that cause Huntington’s disease and Friedreich’s ataxia reduces somatic repeat expansions in patient cells and in mice. Nature Genetics 2025.

44. Saha K: Base editing as a therapeutic strategy for somatic repeat expansion diseases. Nature Genetics 2025.

45. Li J, Lan Z, Liao W, Horner JW, Xu X, Liu J, Yoshihama Y, Jiang S, Shim HS, Slotnik M, et al: Histone demethylase KDM5D upregulation drives sex differences in colon cancer. Nature 2023, 619:632–639.

46. Qin Q, Wang D, Qu Y, Li J, An K, Mao Z, Li J, Xiong Y, Min Z, Xue Z: Enhanced glycolysis-derived lactate promotes microglial activation in Parkinson’s disease via histone lactylation. npj Parkinson’s Disease 2025, 11:3.

47. Gu A, Dao T: Mamba: Linear-time sequence modeling with selective state spaces. arXiv preprint arXiv:231200752 2023.

48. Kokhlikyan N, Miglani V, Martin M, Wang E, Alsallakh B, Reynolds J, Melnikov A, Kliushkina N, Araya C, Yan S: Captum: A unified and generic model interpretability library for pytorch. arXiv preprint arXiv:200907896 2020.

