## supplementary for "Predicting Disease-Specific Histone Modifications and Functional Effects of Non-coding Variants by Leveraging DNA Language Models"

Table S1. The dataset overview of this study

|  | Sample ID | Years | Gender | H3K27ac | H3K4me3 | H3K27me3 |
| --- | --- | --- | --- | --- | --- | --- |
| No Cognitive Impairment | ENCDO250PFZ | 90+ years | Female | ENCFF721ZGP | ENCFF350LHB | ENCFF710FUW |
|  | ENCDO203ASI | 87 years | Male | ENCFF761QUK | ENCFF491IPL | ENCFF556XGB |
|  | ENCDO218FFZ | 90+ years | Female | ENCFF955SXY | ENCFF686AFC | ENCFF864YZX |
|  | ENCDO609ZOG | 79 years | Female | ENCFF289MDK | ENCFF525EJT | ENCFF079QBO |
|  | ENCDO592ZWW | 83 years | Male | ENCFF337DEA | ENCFF344TGW | ENCFF026LAQ |
|  | ENCDO623FPG | 78 years | Male | ENCFF508PQT | ENCFF196MXR | ENCFF321ZZK |
| Mild Cognitive Impairment | ENCDO471EKG | 90+ years | Female | ENCFF672SGH | ENCFF751BEG | ENCFF360LHN |
|  | ENCDO672KST | 90+ years | Female | ENCFF010VMY | ENCFF678JLE | ENCFF169UVC |
|  | ENCDO832DBZ | 83 years | Female | ENCFF872ZJD | ENCFF959JIF | ENCFF218KRJ |
|  | ENCDO877NVF | 90+ years | Male | ENCFF318GXB | ENCFF118NVV | ENCFF784CFJ |
|  | ENCDO697SWU | 89 years | Male | ENCFF384SYD | ENCFF954FED | ENCFF370CQH |
| Cognitive Impairment | ENCDO077CCP | 81 years | Female | ENCFF189UQY | ENCFF774BBN | ENCFF795SLA |
|  | ENCDO359XWR | 90+ years | Female | ENCFF970PFD | ENCFF825JIZ | ENCFF121GIC |
|  | ENCDO448YMQ | 90+ years | Female | ENCFF045ADI | ENCFF700VSF | ENCFF166LAY |
|  | ENCDO845GYA | 86 years | Female | ENCFF283HUT | ENCFF629UTT | ENCFF258VXT |
| Alzheimer's Disease | ENCDO258GJF | 90+ years | Female | ENCFF509IXE | ENCFF285XKT | ENCFF051WDV |
|  | ENCDO997SGX | 86 years | Female | ENCFF283LVU | ENCFF855GAM | ENCFF689TKU |
|  | ENCDO201EUI | 90+ years | Female | ENCFF528AIU | ENCFF653PXI | ENCFF620QQE |
|  | ENCDO853VGZ | 89 years | Female | ENCFF983DQA | ENCFF051DLA | ENCFF276LSC |
| Alzheimer's Disease and Cognitive Impairment | ENCDO940VNT | 87 years | Female | ENCFF024XNY | ENCFF261NKP | ENCFF889LTS |
|  | ENCDO776JEI | 87 years | Male | ENCFF756JDB | ENCFF038YCT | ENCFF736DVV |
|  | ENCDO754RFQ | 90+ years | Female | ENCFF860TAY | ENCFF282ILQ | ENCFF082HRF |

Table S2. Performance comparison of micro-average AUROCs between our model and other state-of-the-art models based on independent tests.

| Models | H3K27ac | H3K4me3 | H3K27me3 |
| --- | --- | --- | --- |
| DeepSEA | 0.7759 | 0.8747 | 0.7729 |
| DeepHistone | 0.7891 | 0.8834 | 0.7766 |
| Expecto | 0.8039 | 0.8979 | 0.7902 |
| Our model | **0.8170** | **0.9142** | **0.8171** |

*The best performance is highlighted in bold.

Table S3. Performance comparison of average AUROCs between our model and other state-of-the-art models based on independent tests.

| Models | H3K27ac | H3K4me3 | H3K27me3 |
| --- | --- | --- | --- |
| DeepSEA | 0.7716(0.0231) | 0.7262(0.0383) | 0.8702(0.0145) |
| DeepHistone | 0.7810(0.0233) | 0.7318(0.0356) | 0.8794(0.0127) |
| Expecto | 0.7967(0.0225) | 0.7507(0.0368) | 0.8928(0.0150) |
| Our model | **0.8104(0.0244)** | **0.7863(0.0307)** | **0.9117(0.0139)** |

*Values are reported as mean (standard deviation) over all models. The best performance is highlighted in bold.

Table S4. Performance comparison of micro-average AUROCs between different variations of our model based on independent tests.

| Models | H3K27ac | H3K4me3 | H3K27me3 |
| --- | --- | --- | --- |
| CNN+BLSTM | 0.8107 | 0.9060 | 0.8051 |
| CNN+MOE | 0.7786 | 0.8830 | 0.7927 |
| LLM+BLSTM | 0.8149 | 0.9117 | 0.8154 |
| LLM+MOE | **0.8170** | **0.9142** | **0.8171** |

*The best performance is highlighted in bold.

Table S5. Performance comparison of average AUROCs between different variations of our model based on independent tests.

| Models | H3K27ac | H3K4me3 | H3K27me3 |
| --- | --- | --- | --- |
| CNN+BLSTM | 0.8039(0.0199) | 0.7714(0.0316) | 0.8979(0.0154) |
| CNN+MOE | 0.7713(0.0222) | 0.7528(0.0368) | 0.8788(0.0216) |
| LLM+BLSTM | 0.7713(0.0222) | 0.7528(0.0368) | 0.8788(0.0216) |
| LLM+MOE | **0.8104(0.0244)** | **0.7863(0.0307)** | **0.9117(0.0139)** |

*Values are reported as mean (standard deviation) over all models. The best performance is highlighted in bold.

Table S6 Performance comparison between different lengths based on independent tests.

| Metrics | Length | H3K27ac | H3K4me3 | H3K27me3 |
| --- | --- | --- | --- | --- |
| AUROC | 512 | 0.7673 | 0.8664 | 0.7684 |
|  | 1024 | 0.7886 | 0.8873 | 0.7825 |
|  | 2048 | 0.8063 | 0.8973 | 0.7962 |
|  | 4096 | **0.8170** | 0.9142 | 0.8171 |
|  | 5120 | 0.8155 | **0.9157** | **0.8248** |
| AUPRC | 512 | 0.5580 | 0.7440 | 0.3863 |
|  | 1024 | 0.5887 | 0.7788 | 0.4106 |
|  | 2048 | 0.6121 | 0.7954 | 0.4361 |
|  | 4096 | **0.6246** | 0.8252 | 0.4599 |
|  | 5120 | 0.6215 | **0.8283** | **0.4723** |

*The best performance is highlighted in bold.

Table S7 KEGG enrichments for differential H3K27ac sites.

| geneSet | description | size | overlap | expect | enrichmentRatio | pValue | FDR |
| --- | --- | --- | --- | --- | --- | --- | --- |
| hsa04720 | Long-term potentiation | 67 | 21 | 8.4176 | 2.4948 | 4.24E-05 | 0.0019 |
| hsa04520 | Adherens junction | 93 | 29 | 11.6842 | 2.4820 | 1.75E-06 | 0.0002 |
| hsa05100 | Bacterial invasion of epithelial cells | 77 | 24 | 9.6740 | 2.4809 | 1.36E-05 | 0.0012 |
| hsa05412 | Arrhythmogenic right ventricular cardiomyopathy | 84 | 25 | 10.5534 | 2.3689 | 2.21E-05 | 0.0013 |
| hsa04730 | Long-term depression | 60 | 17 | 7.5382 | 2.2552 | 0.00083359 | 0.0091 |
| hsa04971 | Gastric acid secretion | 76 | 21 | 9.5484 | 2.1993 | 0.00031235 | 0.0048 |
| hsa04925 | Aldosterone synthesis and secretion | 98 | 26 | 12.3124 | 2.1117 | 0.00013206 | 0.0031 |
| hsa05032 | Morphine addiction | 91 | 24 | 11.4329 | 2.0992 | 0.00026073 | 0.0047 |
| hsa04724 | Glutamatergic synapse | 115 | 30 | 14.4482 | 2.0764 | 5.83E-05 | 0.0019 |
| hsa04211 | Longevity regulating pathway | 89 | 23 | 11.1816 | 2.0569 | 0.00047732 | 0.0070 |
| hsa05410 | Hypertrophic cardiomyopathy | 97 | 25 | 12.1867 | 2.0514 | 0.00028846 | 0.0047 |
| hsa04070 | Phosphatidylinositol signaling system | 97 | 25 | 12.1867 | 2.0514 | 0.00028846 | 0.0047 |
| hsa04725 | Cholinergic synapse | 113 | 29 | 14.1969 | 2.0427 | 0.00010571 | 0.0029 |
| hsa04152 | AMPK signaling pathway | 121 | 31 | 15.2020 | 2.0392 | 6.41E-05 | 0.0019 |
| hsa04911 | Insulin secretion | 86 | 22 | 10.8047 | 2.0361 | 0.00072969 | 0.0091 |
| hsa04728 | Dopaminergic synapse | 132 | 33 | 16.5840 | 1.9899 | 6.37E-05 | 0.0019 |
| hsa04071 | Sphingolipid signaling pathway | 121 | 30 | 15.2020 | 1.9734 | 0.00015861 | 0.0035 |
| hsa04666 | Fc gamma R-mediated phagocytosis | 97 | 24 | 12.1867 | 1.9694 | 0.00071952 | 0.0091 |
| hsa04261 | Adrenergic signaling in cardiomyocytes | 154 | 38 | 19.3480 | 1.9640 | 2.50E-05 | 0.0013 |
| hsa04015 | Rap1 signaling pathway | 210 | 51 | 26.3836 | 1.9330 | 1.79E-06 | 0.0002 |
| hsa05414 | Dilated cardiomyopathy | 103 | 25 | 12.9405 | 1.9319 | 0.00076272 | 0.0091 |
| hsa04144 | Endocytosis | 250 | 59 | 31.4091 | 1.8784 | 7.79E-07 | 0.0002 |
| hsa04919 | Thyroid hormone signaling pathway | 121 | 28 | 15.2020 | 1.8419 | 0.00085565 | 0.0091 |
| hsa04072 | Phospholipase D signaling pathway | 148 | 34 | 18.5942 | 1.8285 | 0.0002907 | 0.0047 |
| hsa04514 | Cell adhesion molecules | 158 | 36 | 19.8505 | 1.8136 | 0.00023188 | 0.0047 |
| hsa04360 | Axon guidance | 182 | 41 | 22.8658 | 1.7931 | 0.00011468 | 0.0029 |
| hsa04371 | Apelin signaling pathway | 139 | 31 | 17.4634 | 1.7751 | 0.00089986 | 0.0092 |
| hsa04810 | Regulation of actin cytoskeleton | 229 | 51 | 28.7707 | 1.7726 | 2.44E-05 | 0.0013 |
| hsa04218 | Cellular senescence | 156 | 34 | 19.5993 | 1.7348 | 0.00080413 | 0.0091 |
| hsa04148 | Efferocytosis | 156 | 34 | 19.5993 | 1.7348 | 0.00080413 | 0.0091 |
| hsa05163 | Human cytomegalovirus infection | 225 | 49 | 28.2682 | 1.7334 | 6.44E-05 | 0.0019 |
| hsa04022 | cGMP-PKG signaling pathway | 167 | 36 | 20.9813 | 1.7158 | 0.0007098 | 0.0091 |
| hsa04530 | Tight junction | 170 | 36 | 21.3582 | 1.6855 | 0.00100033 | 0.0098 |
| hsa05205 | Proteoglycans in cancer | 205 | 42 | 25.7554 | 1.6307 | 0.00080824 | 0.0091 |
| hsa04024 | cAMP signaling pathway | 225 | 45 | 28.2682 | 1.5919 | 0.0009125 | 0.0092 |

Table S8 KEGG enrichments for differential H3K4me3 sites.

| geneSet | description | size | overlap | expect | enrichmentRatio | pValue | FDR |
| --- | --- | --- | --- | --- | --- | --- | --- |
| hsa04012 | ErbB signaling pathway | 85 | 21 | 8.3584 | 2.5124 | 0.0001 | 0.0062 |
| hsa05412 | Arrhythmogenic right ventricular cardiomyopathy | 84 | 19 | 8.2601 | 2.3002 | 0.0004 | 0.0205 |
| hsa04330 | Notch signaling pathway | 62 | 14 | 6.0967 | 2.2963 | 0.0023 | 0.0326 |
| hsa05100 | Bacterial invasion of epithelial cells | 77 | 17 | 7.5717 | 2.2452 | 0.0011 | 0.0273 |
| hsa05032 | Morphine addiction | 91 | 20 | 8.9484 | 2.2350 | 0.0004 | 0.0205 |
| hsa04927 | Cortisol synthesis and secretion | 65 | 14 | 6.3917 | 2.1903 | 0.0037 | 0.0443 |
| hsa04520 | Adherens junction | 93 | 20 | 9.1451 | 2.1870 | 0.0006 | 0.0205 |
| hsa04911 | Insulin secretion | 86 | 18 | 8.4567 | 2.1285 | 0.0015 | 0.0277 |
| hsa05223 | Non-small cell lung cancer | 72 | 15 | 7.0801 | 2.1186 | 0.0038 | 0.0443 |
| hsa04921 | Oxytocin signaling pathway | 154 | 32 | 15.1435 | 2.1131 | 0.0000 | 0.0055 |
| hsa04713 | Circadian entrainment | 97 | 20 | 9.5384 | 2.0968 | 0.0010 | 0.0273 |
| hsa00562 | Inositol phosphate metabolism | 73 | 15 | 7.1784 | 2.0896 | 0.0043 | 0.0483 |
| hsa04725 | Cholinergic synapse | 113 | 23 | 11.1118 | 2.0699 | 0.0005 | 0.0205 |
| hsa04360 | Axon guidance | 182 | 37 | 17.8968 | 2.0674 | 0.0000 | 0.0045 |
| hsa05414 | Dilated cardiomyopathy | 103 | 20 | 10.1284 | 1.9746 | 0.0022 | 0.0323 |
| hsa04270 | Vascular smooth muscle contraction | 134 | 26 | 13.1768 | 1.9732 | 0.0005 | 0.0205 |
| hsa04925 | Aldosterone synthesis and secretion | 98 | 19 | 9.6367 | 1.9716 | 0.0029 | 0.0372 |
| hsa04935 | Growth hormone synthesis, secretion and action | 120 | 23 | 11.8001 | 1.9491 | 0.0013 | 0.0277 |
| hsa04152 | AMPK signaling pathway | 121 | 23 | 11.8984 | 1.9330 | 0.0014 | 0.0277 |
| hsa04919 | Thyroid hormone signaling pathway | 121 | 23 | 11.8984 | 1.9330 | 0.0014 | 0.0277 |
| hsa04928 | Parathyroid hormone synthesis, secretion and action | 106 | 20 | 10.4234 | 1.9188 | 0.0031 | 0.0393 |
| hsa04261 | Adrenergic signaling in cardiomyocytes | 154 | 29 | 15.1435 | 1.9150 | 0.0004 | 0.0205 |
| hsa04022 | cGMP-PKG signaling pathway | 167 | 31 | 16.4218 | 1.8877 | 0.0004 | 0.0205 |
| hsa04072 | Phospholipase D signaling pathway | 148 | 27 | 14.5534 | 1.8552 | 0.0011 | 0.0273 |
| hsa04611 | Platelet activation | 124 | 22 | 12.1934 | 1.8043 | 0.0044 | 0.0483 |
| hsa04728 | Dopaminergic synapse | 132 | 23 | 12.9801 | 1.7719 | 0.0046 | 0.0488 |
| hsa04510 | Focal adhesion | 203 | 35 | 19.9618 | 1.7533 | 0.0007 | 0.0211 |
| hsa04530 | Tight junction | 170 | 29 | 16.7168 | 1.7348 | 0.0022 | 0.0323 |
| hsa04140 | Autophagy | 165 | 28 | 16.2251 | 1.7257 | 0.0028 | 0.0372 |
| hsa05130 | Pathogenic Escherichia coli infection | 198 | 33 | 19.4702 | 1.6949 | 0.0017 | 0.0281 |
| hsa04024 | cAMP signaling pathway | 225 | 37 | 22.1252 | 1.6723 | 0.0012 | 0.0273 |
| hsa04020 | Calcium signaling pathway | 253 | 40 | 24.8785 | 1.6078 | 0.0016 | 0.0281 |

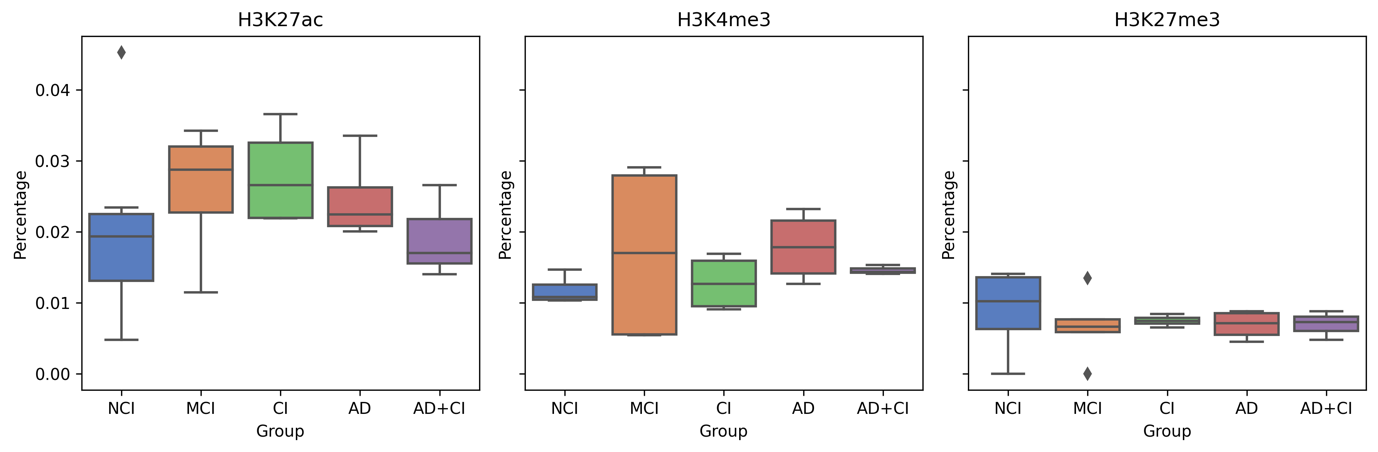

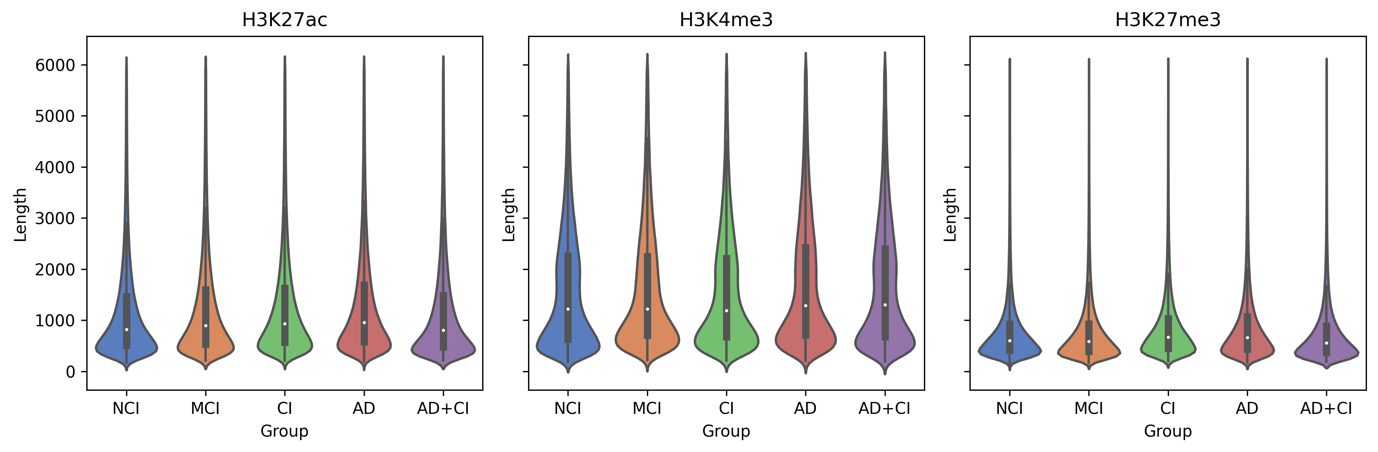

Figure S1 Distribution of the length of the histone modification sites

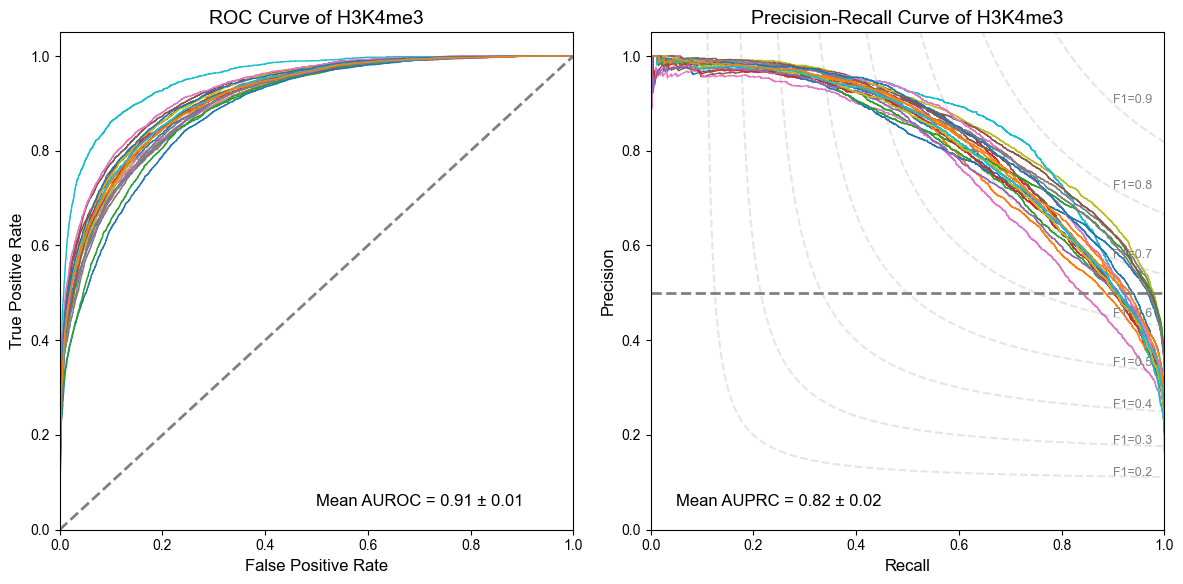

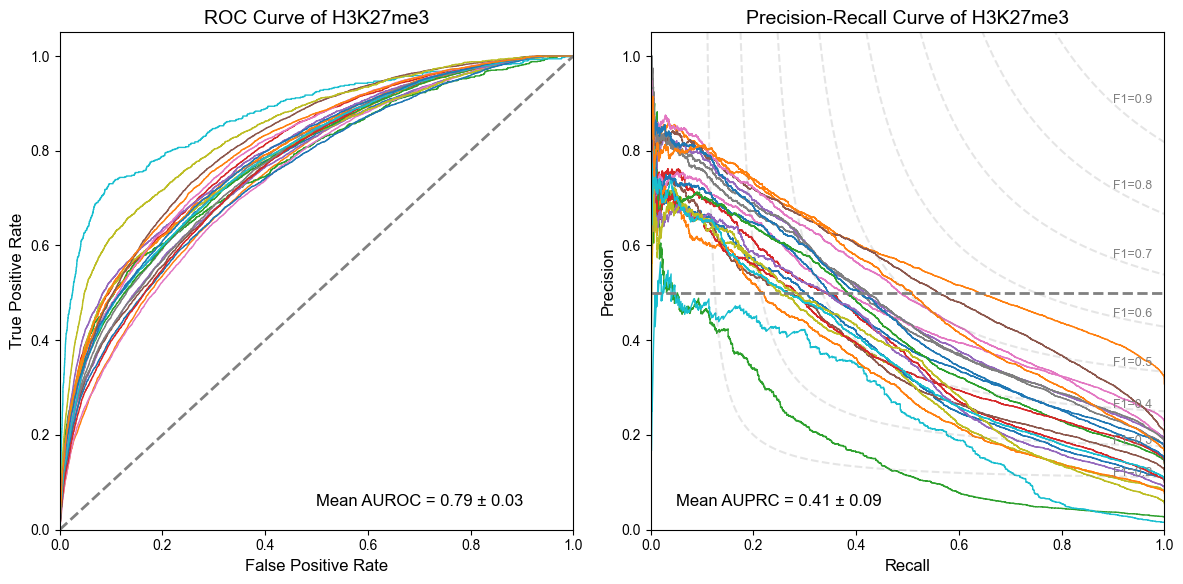

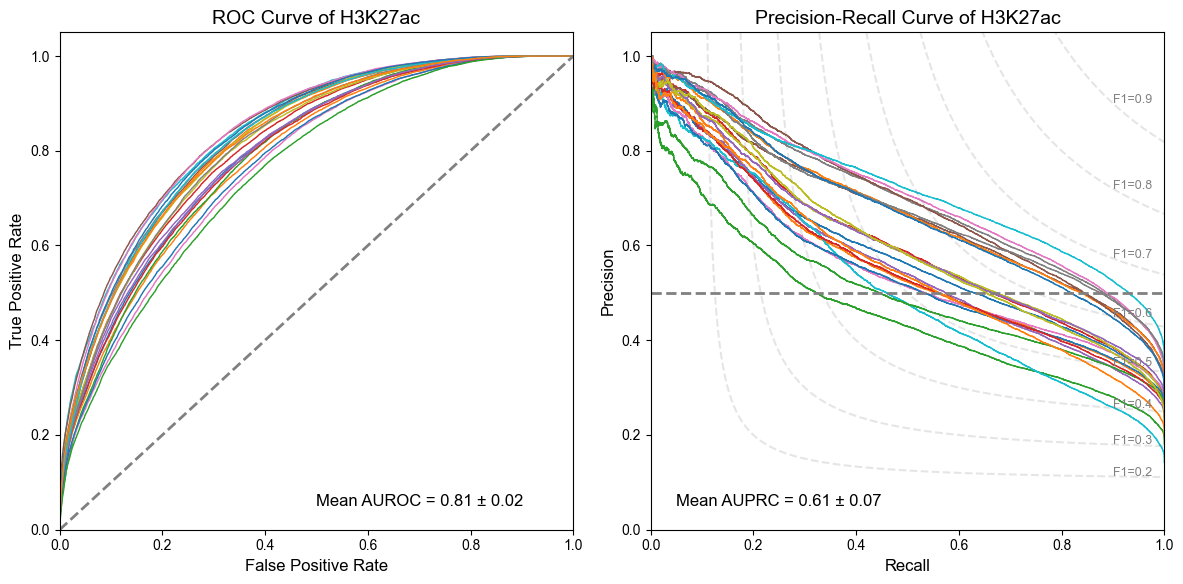

Figure S2. AUROCs and AUPRCs of our model.

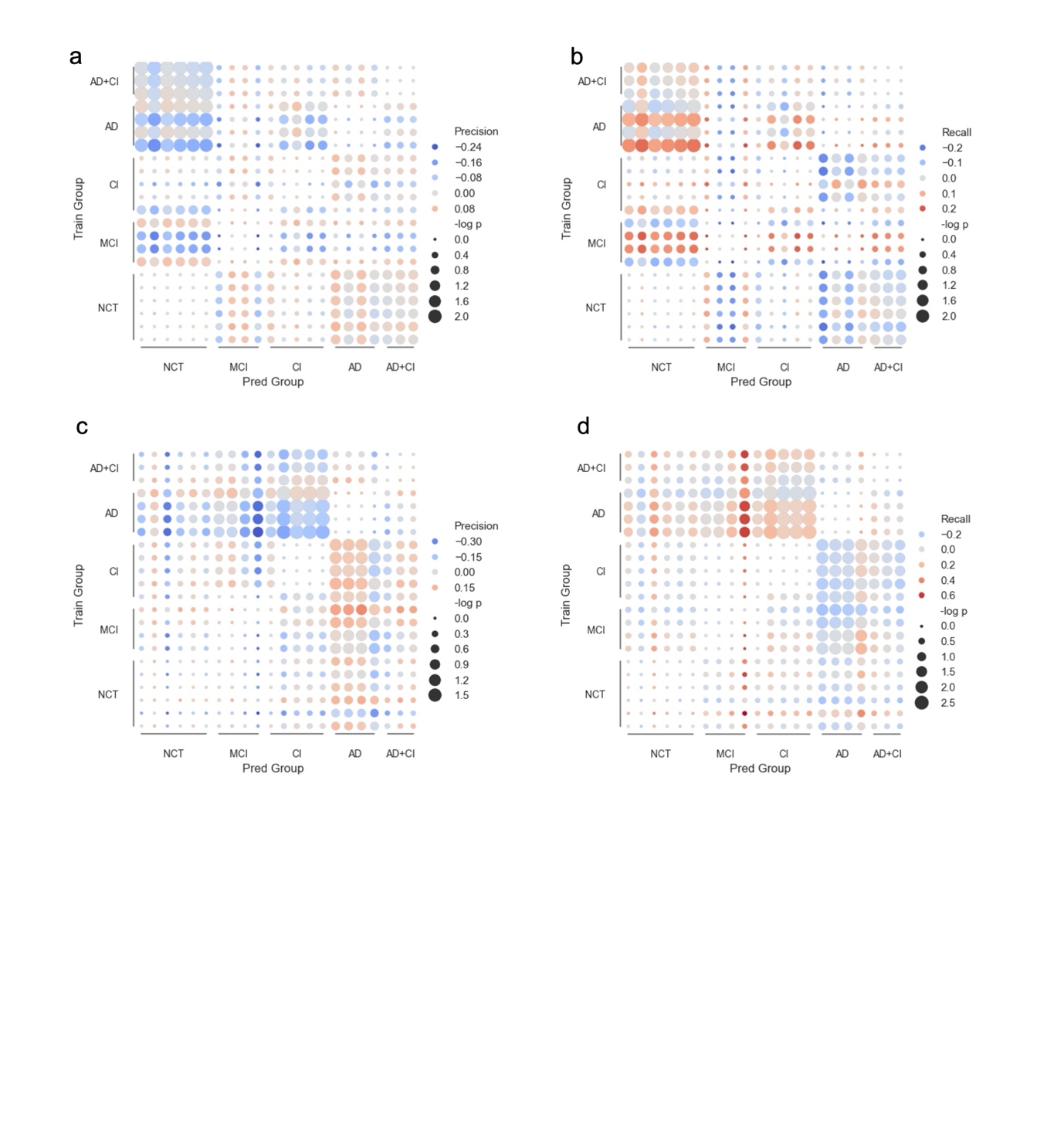

Figure S3. The cross-disease analysis for H3K4me3(a,b) and H3K27ac (c,d) models. (a,c) Precision and (b,d) recall metrics across disease groups, where dot size represents ${-log}_{10} P$ value and color gradient indicates performance deviation in cross-disease analysis.
